# Deregulated Myt3 translation predisposes islet β-cells to dysfunction under obesity-induced metabolic stress

**DOI:** 10.1101/2025.05.11.653323

**Authors:** Ruiying Hu, Yu Wang, Mahircan Yagan, Yanwen Xu, Alan J. Simmons, Ken S. Lau, Qi Liu, Guoqiang Gu

## Abstract

In response to obesity-related metabolic stress, islet β-cells adapt (or compensate) by increasing their secretory function and mass. Yet, for unknown reasons, this compensation is reversed in some individuals at some point to induce β-cell failure and overt type 2 diabetes (T2D). We have previously shown that transcription factor Myt3 (St18) and its paralogs, Myt1 and Myt2, prevent β-cell failure. Myt3 was induced at post-transcriptional levels by obesity-related stress in both mouse and human β cells and its downregulation accompanied β-cell dysfunction during T2D development. Single-nucleotide polymorphisms in *MYT3* were associated with an increased risk of developing human diabetes. We now demonstrate that Myt3 translation is regulated by an upstream open-reading frame that overlaps with the main Myt3 open-reading frame in mice. Disrupting this overlap enhances Myt3 translation in mouse β cells without metabolic stress but decreases it under high-fat-diet challenges. Consequently, this deregulation results in β-cell dysfunction and glucose intolerance in mice, accompanied by compromised expression of several β-cell function genes. These findings suggest that stress-induced Myt3 translation is part of the compensation mechanism that prevents β-cell failure.

## Introduction

Type 2 diabetes (T2D) arises when endocrine islet β cells cannot secrete enough insulin to regulate blood glucose homeostasis (1,2). This disease usually starts with obesity-related insulin resistance. In response, β cells increase their mass and secretory function to boost insulin output. This adaptation (or compensation) can maintain lifetime glucose homeostasis in most obese subjects (3). Yet in ∼20% of these people, adaptation stops after some time, and β-cell failure follows in the forms of β-cell loss of identity, death, and/or dysfunction (i.e., β-cell failure), leading to overt diabetes. The mechanisms that cause the transition from compensation to failure are unknown but are thought to be related to workload-related stress response (4).

Producing and driving insulin secretion are metabolically stressful processes. During insulin biosynthesis, a large amount of unfolded proinsulin can be made in the ER (5). If not removed, the unfolded proteins will decimate the ER function, inducing β-cell failure. To stimulate insulin secretion, high levels of glucose metabolism are needed to increase the ATP/ADP ratio that drives subsequent membrane depolarization, Ca^2+^ influx, and secretion. Glucose metabolism co-produces <u>r</u>eactive <u>o</u>xygen <u>s</u>pecies (ROS), which at low levels enhance GSIS but at high levels induce cell dysfunction and/or death (6–8). Therefore, β cells activate the unfolded protein response (UPR) and oxidative stress response (OSR) to clear these toxic products (9). During UPR, cells also induce chaperones, chaperonins, and proteases (involved in ER-aided degradation) via IRE1α- and Atf6-mediated RNA processing and transcription (5,10). They also reduce overall protein translation while inducing the selective production of specific stress effector molecules such as Atf4 (11). This selective protein translation is made possible by the PERK-eIF2α axis that activates the translation of mRNAs with particular features in their 5’ end, including short upstream open-reading-frames (uORF) like that in Atf4 (11). During OSR, the transcription factor Nurf2 is activated to induce enzymes for the removal of ROS (12,13). The overall result of these responses is a reduction in unfolded proteins and ROSs, whose deregulation leads to β-cell dysfunction and diabetes (6).

An overly activated stress response, however, can repress key β-cell factors (14,15) while also activating some downstream proapoptotic effectors such as Atf4 target genes CHOP and Bid (2,5,16,17). Thus, mechanisms that guard against the overactivation of stress response likely play an essential role in preventing the transition from β-cell adaptation to failure (18).

We have recently identified a family of myelin transcription factors, Myt1, 2, and 3, that prevent β-cell failure by repressing a few stress response genes (19,20). Amongst these family members, Myt3 is particularly intriguing. Two *MYT3* single-nucleotide polymorphisms are associated with the risk of human diabetes (21,22). Its transcript levels anticorrelate with the secretory function of primary human β cells; its protein levels are induced by obesity-related metabolic stress in functional rodent and human donor β cells, yet are decreased in failing primary mouse and human β cells (20). Here, we examine how deregulating Myt3 translation impacts β-cell adaptation in mice.

## Results

### Myt3 is required for robust glucose-induced insulin secretion from β cells

Co-inactivation of all three *Myt* genes in pancreatic cells using a transgenic mouse *Pdx1^Cre^* led to overt diabetes in young adult mice, whereas inactivating *Myt3* alone did not (20). We therefore tested whether the *Myt3^F/F^; Pdx1^Cre^* mice exhibited more subtle defects in glucose homeostasis and β-cell function, such as glucose clearance or glucose-stimulated insulin secretion (GSIS) in isolated islets.

Over 95% of islet cells have lost Myt3 production in *Myt3^F/F^; Pdx1^Cre^* mice (Fig. 1A, B). In young adults (∼6 weeks) after birth, neither male nor female mutants showed defective glucose clearance (Fig. S1A), accompanied by normal glucose-stimulated insulin secretion (GSIS) in isolated islets (Fig. S1B). By ∼4-5 months after birth, both males and females showed defective glucose clearance (Fig. S1C, D and Fig. 1C). Consistent with this physiological defect, isolated islets from these mice showed compromised glucose-induced insulin secretion (GSIS) (Fig. 1D). These results established the roles of Myt3 in β cells, leading us to explore how deregulating obesity-induced Myt3 up-regulation impacted β-cell dysfunction.

**Fig. 1.**
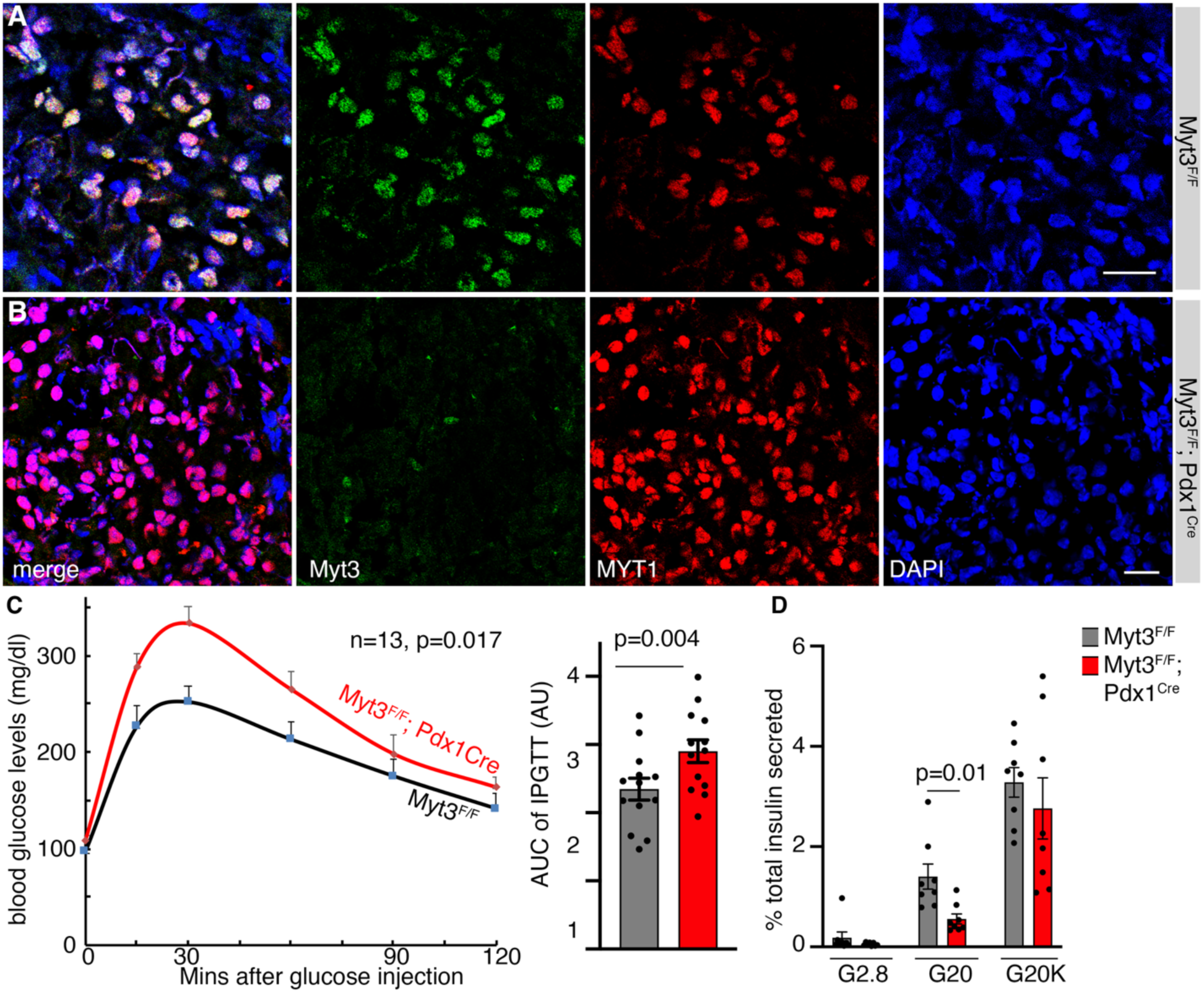
Myt3 is required for β-cell GSIS in adult mice. (A, B) Myt3 inactivation efficacy by *Pdx1^Cre^*. Merged images and single-channel images were displayed. Myt1 staining was included to highlight the co-expression of Myt1 and MYT3. DAPI was used to note the location of nuclei. Bars, 20 μm. (C) IPGTT results in mice ∼4-5 months after birth, with both scatter-plot and Area-Under-Curve (AUC) shown. For both the control and mutant groups, 6 females and 7 males were included. The P-values are from repeated-measure ANOVA (line-graph) and a t-test with a two-tailed type 2 error (AUC). (D) GSIS assays of isolated islets (5-6 months old). Both male and female islets were included. Each dot represents one independent assay. Assays were done on two days. Each day, four GSIS assays were done for control (from two males + two females) and mutant samples (each with two males + two females). The P-value is from a t-test with a two-tailed type 2 error.

### Myt3 mRNAs have features that allow their stress-regulated translation

Several stress response effectors are known to be induced at translational levels because of some special features in the 5’ non-coding regions. One of these features is the presence of uORFs, short ORFs that either terminate before or overlap with the main ORF of functional proteins. The uORFs normally inhibit translation. Yet under cellular stress when eIF2α is phosphorylated, the presence of uORFs improve the chance of recruiting the ribosomal initiation complex to the start codon of the main ORF for higher translational efficacy (11). For this reason, we analyzed whether uORFs exist in the 5’ end of *Myt3* mRNA.

To identify the primary *Myt3* transcripts, we utilized our RNA-seq results from purified adult mouse cells (23). These data were generated using random priming so that aligning the sequenced cDNA fragments against the *Myt3* genomic sequence will show the potential transcriptional starting site (TSS) and splicing patterns (Fig. S2A). This analysis identified 25 exons with detectable reads from the *Myt3* locus (Fig. 2A). These include one TSS and 5 exons involved in alternative splicing. Exons 9-25 were found in >95% of all cDNAs, while exons E3, E4, and E6 were found in less than 5% of cDNAs. This splicing pattern allowed us to identify two primary *Myt3* transcripts (T1 and T2), which were verified via RT-PCR using oligos in exon 1 and exon 9, followed by sequencing (Fig. 2A). By inspecting the sequences upstream of the main ORF for Myt3 protein, we found that both transcripts have three uORFs. In the T1 transcript, none overlaps with the main Myt3 ORF (Fig. 2B). In contrast, the most distal uORF in the T2 transcript overlaps with the main ORF (Fig. 2B), suggesting that T2 translation could be regulated by eIF2α phosphorylation like Atf4, which we tested using a bicistronic reporter (Fig. 2C).

**Fig. 2.**
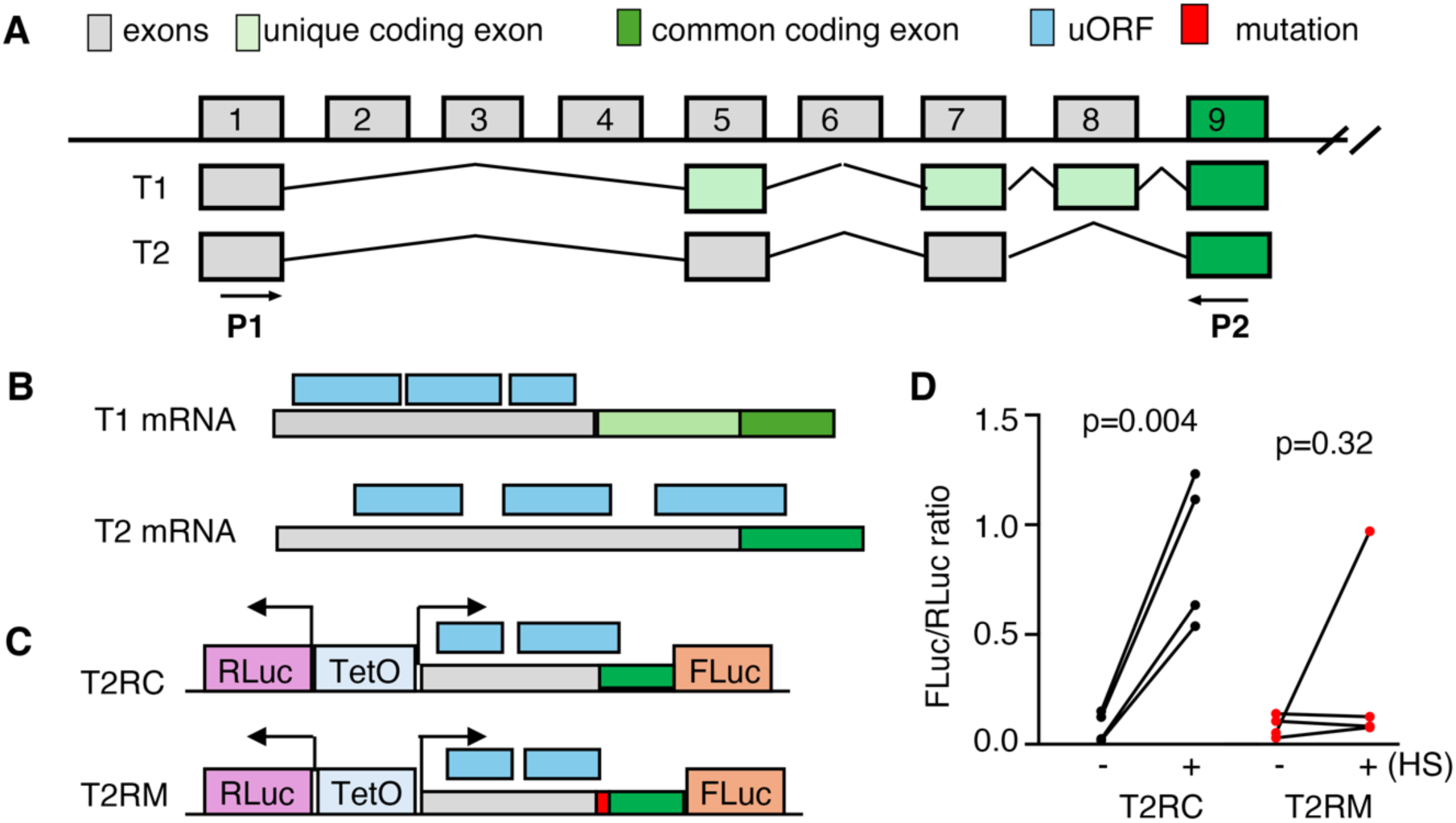
A primary Myt3 transcript has an mRNA element allowing translational control by stress response. (A) Myt3 exons with detectable expression in adult mouse β cells. Only the first 9 of the 25 exons are shown (not in scale). (B) The presence of uORFs in the 5’end of the two primary Myt3 transcripts. (C) Reporter constructs that are used to test stress-induced translation. Note that RLuc and FLuc transcription are all regulated by a TeoO promoter/enhancer, ensuring a constant ratio of their transcription. Also note that in the T2RM construct, a 12 bp sequence (5’-AGCTTTAATGAA-3’) was inserted to disrupt the overlapping uORF. (D) Reporter assay results with or without ER stress response induced by a three-hour heat shock at 42-43 °C. Presented are the ratios between FLuc/RLuc. Each dot represents an independent experiment, each having 2-3 technical duplicates. P-values are from paired t-tests with two-tailed type two errors.

A bi-directional transcription promoter was utilized to drive the simultaneous transcription of Renilla luciferase (RLuc) and firefly luciferase (FLuc) using a TetO-enhancer. In the presence of rTTA, the R-Luc and F-Luc mRNAs were expected to be transcribed at a consistent ratio, with high transcription levels under Dox and low transcription levels without Dox. Thus, their luciferase activity change will be dictated by the translational efficacy of each mRNA. The 5’end of the T2 transcript, together with the expected translation initiation codon of Myt3, was fused in-frame with the F-Luc coding region (Fig. 2C). We found that this sequence can significantly improve the translation of FLuc in the presence of stress, readily induced by a brief heat shock (24).

To test whether the overlapping uORF is required for stress-induced FLuc upregulation, we disrupted this overlap by inserting a 12-bp sequence within the uORF. The inserted sequence was expected to terminate the uORF before reaching the main Myt3 ORF while not affecting the translation from the T1 transcript. This mutation effectively eliminated heat shock-induced Fluc translational increase (Fig. 2C, D), suggesting that Myt1 translation is subject to uORF regulation. These combined findings led us to test whether this particular uORF can regulate Myt3 translation *in vivo* and whether it plays essential roles in β-cell function.

### Eliminating the overlap between the uORF and the primary ORF deregulated Myt3 levels in β cells

A mouse line carrying the desired 12-bp insertion mutation was obtained via CRISPR-CAS9-based editing and verified by DNA fragment digestion and sequencing (Fig. 3A, B). It was back-crossed with wild-type C57Bl/6j mice for four rounds. All the remaining studies were performed using this line, designated as *Myt3^IM^* (or *Myt3* with an insertion mutation).

**Fig. 3.**
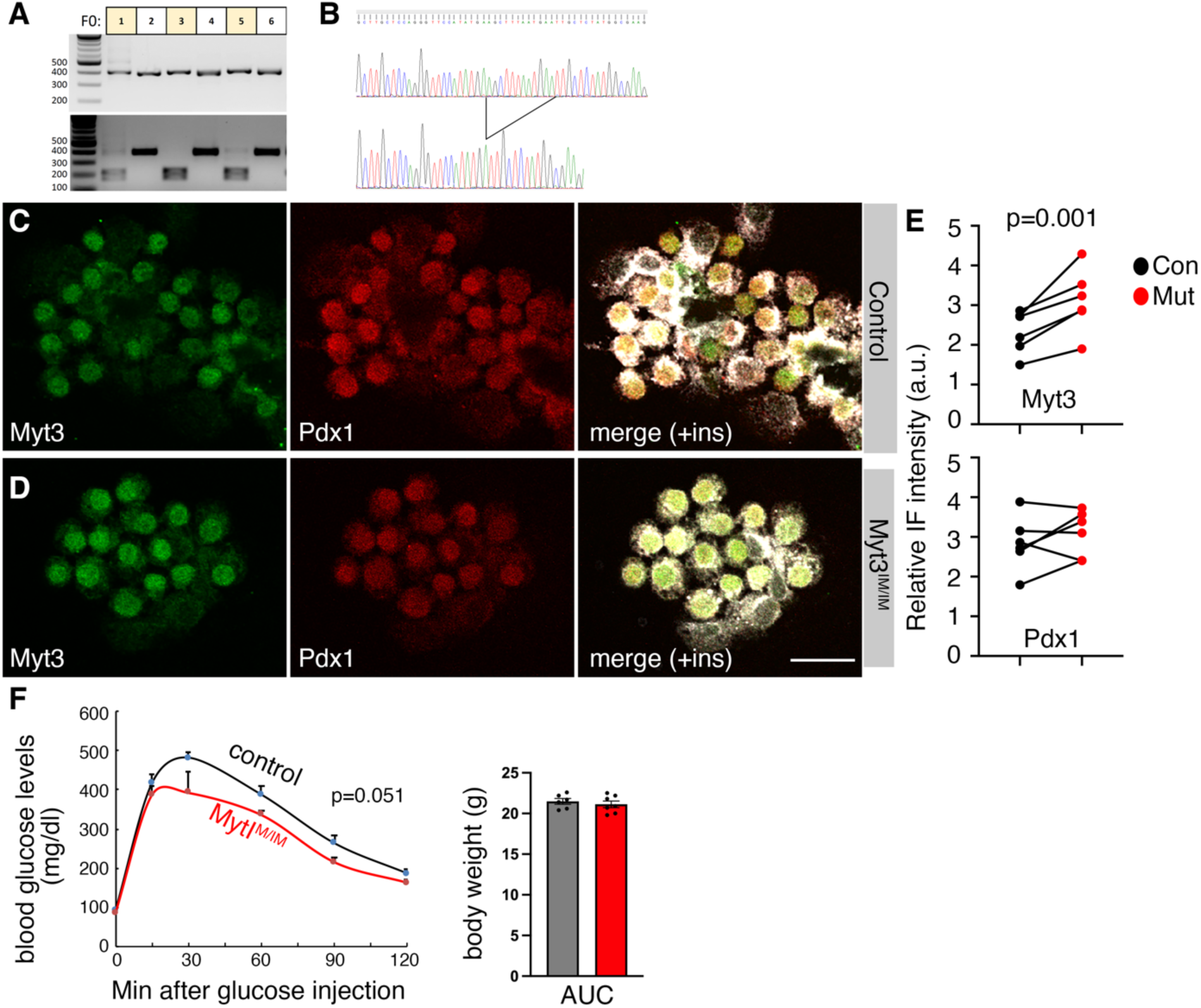
An insertion mutation deregulates Myt3 translation in mouse β cells. (A, B) Verification of the *Myt3^IM^* allele via PCR fragment digestion (A) and sequencing (B). In A, each lane represents one PCR fragment from one animal (top), spanning the mutated site that introduces a new Hind III restriction site that can be tested via digestion (bottom). (C-E) Typical IF staining in isolated islet cells from three-week-old mice. Shown IF panels are average projections from z-stacked images, with single channels and merges of Myt3 and Pdx1 staining. The quantification data in E are relative Myt3 and Pdx1 levels. Each dot represents an average of one mouse (three males and three females). The P-value is from a paired t-test, with a two-tailed type 2 error. (F) IPGTT test of 8-week-old male mice. Six controls and seven mutants were tested. P-value is from two-way ANOVA, repeated measure. (G) Animal body weight at 8 weeks after birth.

Around weaning, β cells from *Myt3^IM/IM^* mice had significantly higher levels of Myt3 protein without significantly changing the levels of Pdx1 (Fig. 3C-E). Additionally, a strong trend of improved glucose clearance is observed in young males (Fig. 3F), without affecting their body weight (Fig. 3G). These results are consistent with the translational reduction effect of the uORFs and the positive role of Myt3 for β-cell secretory function. We next used this mouse line to examine the biological consequences of Myt3 translational deregulation under obesity-related stress.

### *Myt3^IM/IM^* mice have compromised glucose clearance under high-fat diet treatment

Control and *Myt3^IM/IM^* mice were fed a high-fat diet (HFD) for 3-5 months starting from ∼5 weeks after birth. We focused on male mice from now on because a pilot analysis showed that a three-month HFD feeding had no significant effect on glucose tolerance in female mice but compromised glucose clearance in males (Fig. 4A). Corresponding to this observation, islets of HFD-fed mice had significantly blunted insulin secretion (Fig. 4B). There were also compromised Myt3 and Pdx1 production in mutant islet cells (Fig. 4C-E). The proliferation of the β cells in these HFD-treated mutants remained the same in control and mutant mice (Fig. S3A-C). In addition, β-cell apoptosis was undetectable as well (Fig. S3D, E). These findings suggest that deregulated Myt3 production under obesity causes β-cell dysfunction.

**Fig. 4.**
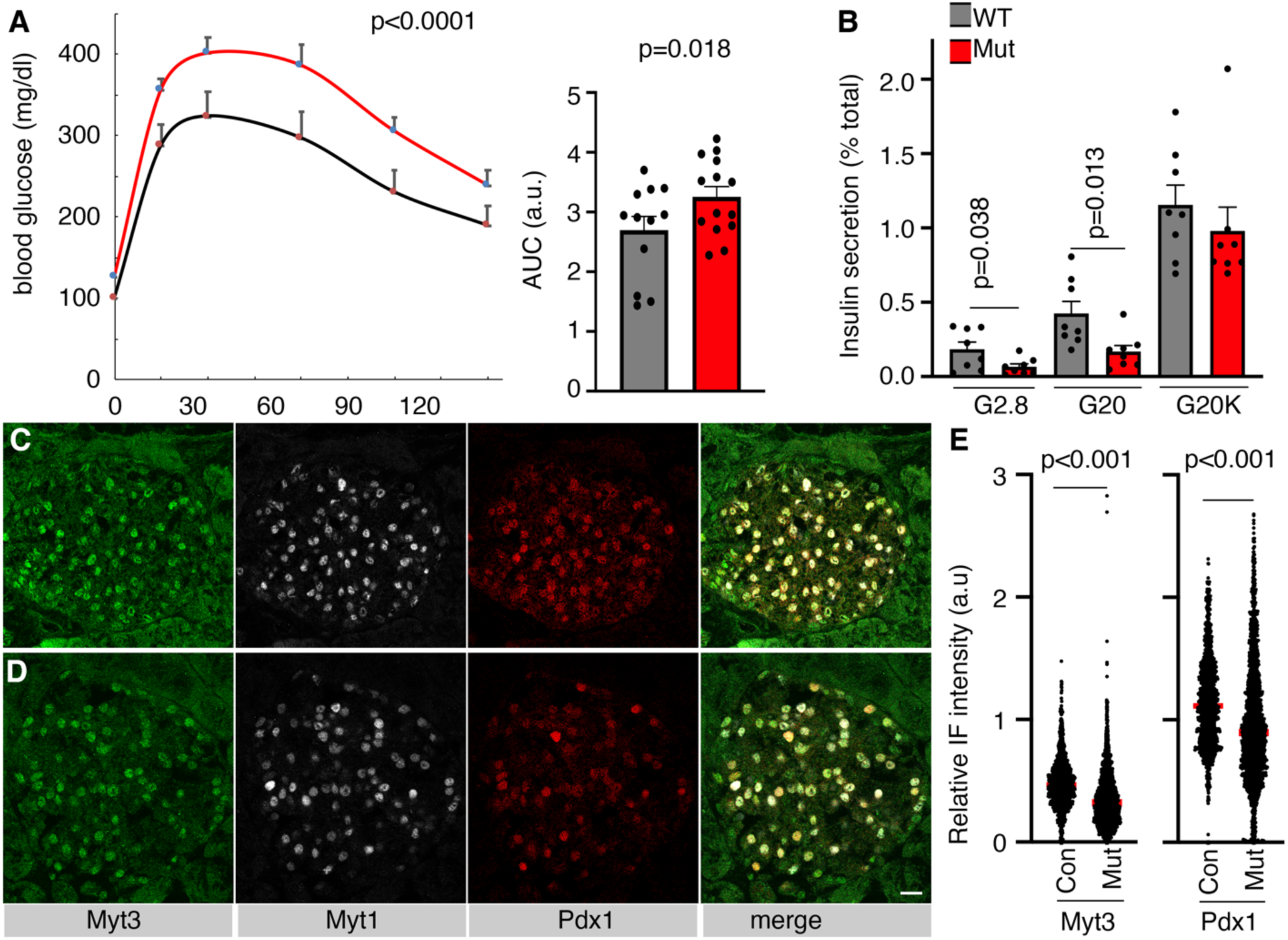
*Myt3^IM/IM^* mice have compromised glucose clearance under HFD treatment. (A) IPGTT results of control and mutant mice after HFD feeding for ∼3-5 months, shown with scatter-plot and AUC. (B) Insulin secretion from islets after 5 months of HFD feeding. (C, D) Myt3 and Pdx1 IF staining in control (C) and mutant (D) pancreatic sections. Bar = 20 μm. (E) Relative Myt3 and Pdx1 levels in the nuclei of control and mutant β cells after 5-month HFD feeding. Each dot represents one cell, with signals integrated from z-stacked images. Three batches of animals (each consisting of 2 mice) were used. The p-values were from a t-test, with two-tailed type 2 errors.

### Myt3 deregulation compromises genes required for β-cell function

We utilized scRNA-seq to identify the altered genes in β-cells in *Myt3^IM/IM^* mice after HFD challenge. From two independent assays of both control and mutant samples, we observed all expected cell types, including the endocrine islet cells (with 8 β-cell subsets) and non-endocrine cells (Fig. 5A). Cell clustering based on cell genotypes showed clear separation of control and mutant cells (Fig. 5B, C), suggesting the profound effect of the introduced *Myt3* mutation. Note that we detected several β-cell subtypes in both control and mutant samples (Fig. 5A-C). Yet β-1, β-2, β-6, and β-10 are mainly detected in the control samples, whereas β-0, β-4, β-7, and β-9 are in mutants. The underlying reason for this observation is unknown and has not been further pursued.

**Fig. 5.**
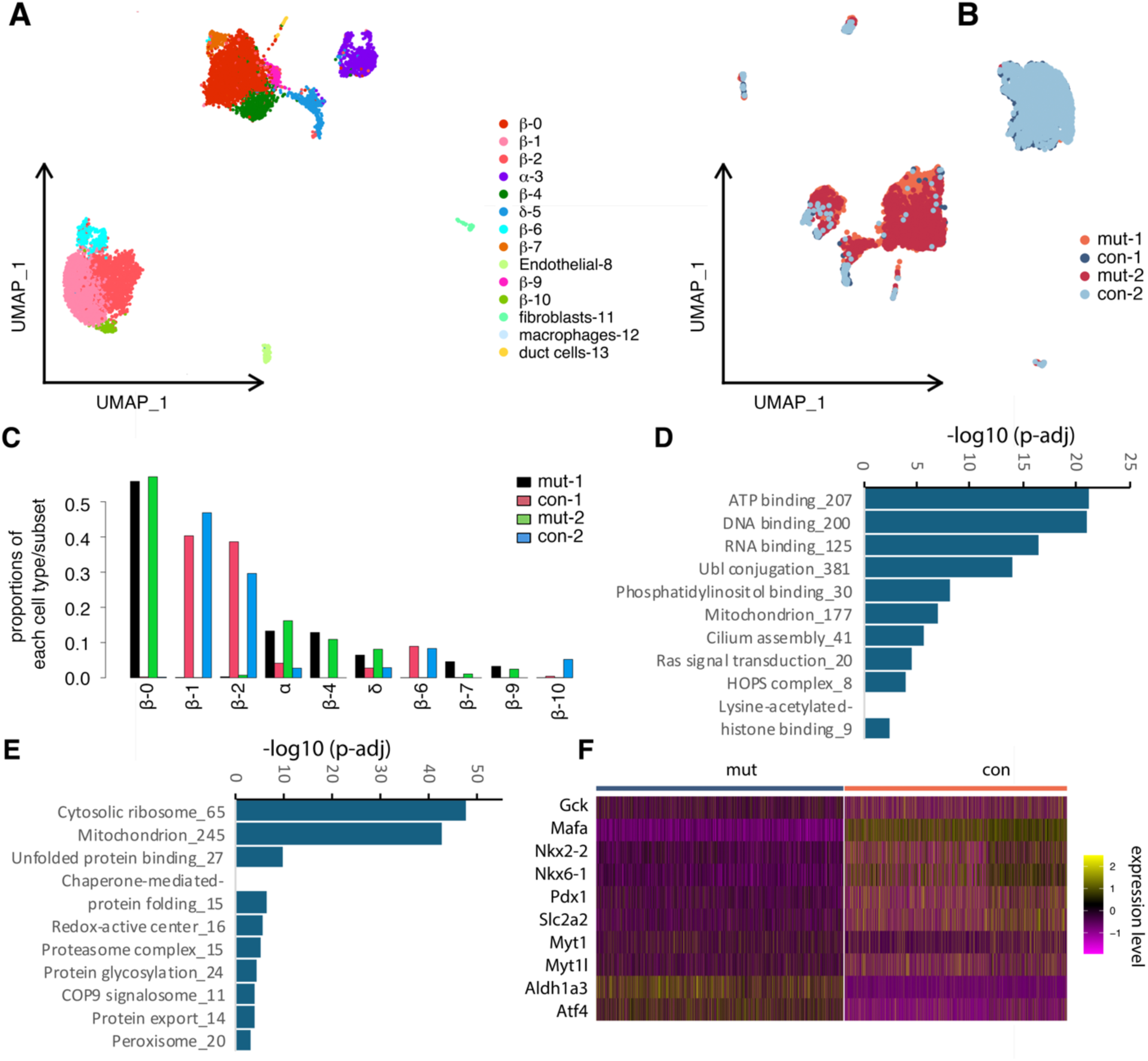
Myt3 deregulation compromised β -cell function genes. (A) UMAP of all cells identified in control and mutant islets clustered based on features of different cell types. (B) UMAP of all cells, clustered based on their sample genotype. The two duplicates were marked using different colors. (D, F) Processes that are down- or up-regulated in mutant β cells after 5-month HFD treatment. (F) Expression of several genes in control and mutant β cells.

By comparing the gene expression differences between the control (3,819 cells) and mutant (3,434) β cells, we identified 3,511 differentially expressed genes (DEGs) (Table S1). Gene ontogeny analysis showed that the downregulated genes regulated processes such as ATP binding, Ubl conjugation, mitochondrion, cilium assembly, Ras signaling, histone modification, etc (Fig. 5D). In contrast, the upregulated genes regulated cytosolic ribosome translation, mitochondrion, unfolded protein binding, chaperones, proteosome, etc. (Fig. 5E). These altered processes are consistent with the compromised secretory function of the *Myt3^IM/IM^* β cells.

Supervised examination of the gene list further supported the dysfunction of *Myt3^IM/IM^* β cells. We did not detect any changes in *Myt3* expression between the control and mutant cells, consistent with our expectation that the insertion mutation would not interfere with *Myt3* transcription or mRNA stability (Table S1). However, the transcript levels of of several diagnostic β-cell markers, *Gck*, *Mafa*, *Nkx2.2*, *Nkx6.1*, *Pdx1*, and *Slc2a2,* were down-regulated (Fig. 5F). The expression of *Myt1* and *Myt2*, was also decreased (Fig. 5F). In contrast, there were increased expression of several markers associated with β-cell dysfunction, including *Aldh1a3* that is associated with dedifferentiation (25,26) and *Atf4* that have opposing roles in β-cell function under high glucose or high fatty acid context (27).

## Discussion

Islet cells’ survival and robust function depend on a cellular stress response that removes toxic stressor molecules to maintain homeostasis. Both inactivation and overactivation of this response can cause β-cell dysfunction and/or death, underscoring the importance of tightly regulating the levels of stress response (9). Here, we investigate a novel mechanism by which MYT3 is regulated at the translational level in response to obesity-related stress. We suggest that Myt3 serves as a rheostat to guard against stress response overactivation, i.e., the transition of β-cell adaptation to failure during T2D development.

Myt3 is a zinc finger transcription factor that is highly expressed in specific neuronal and endocrine islet cells (19). We reported that the co-inactivation of *Myt3* with *Myt1* and *Myt2* compromised β-cell proliferation, survival, and function by de-repressing several stress response effector genes, including Atf4 and heat shock proteins (20). In primary human islets, its knockdown compromised β-cell secretion under normal physiological conditions but resulted in cell death under obesity-related stress (28). The Myt protein levels were upregulated in both mouse and human β cells under obesity-related stress, while their downregulation accompanies β-cell dysfunction in T2D development (20). By completely inactivating *Myt3* in all pancreatic cells, we established an essential role of *Myt3* in mouse-cell insulin secretion in late adult mice. We further demonstrated that translational control of Myt3 is, at least in part, mediated by a uORF that overlaps with the main Myt3 ORF. Disrupting the overlap of this uORF with the Myt3 coding ORF resulted in a slight (∼30%) but significant increase in Myt3 protein levels under normal physiological conditions, consistent with the notion that overlapping uORFs can normally repress protein translation. In contrast, this enhanced protein translation was reversed under obesity-related stress, which causes β-cell dysfunction. Thus, our combined findings underscore the importance of translational control on the Myt3 protein levels during stress response.

Under regular feeding conditions, when mice could maintain normal glucose homeostasis, the increased Myt3 production showed a trend of improving the glucose-clearing capacity in male mice, consistent with the positive roles of this gene in β-cell function. Along a similar line, their reduction, especially under obesity-related metabolic stress, rendered β cells vulnerable to failure. This finding suggests that the lowered Myt3 levels cannot be compensated by its two paralogs, *Myt1* and *Myt2*. We detected decreased transcript levels of *Myt1* and *Myt2* when Myt3 levels were down-regulated. We do not know how *Myt1* and *Myt2* are downregulated and how much their downregulation contributes to the observed -cell defects, whose dosage reduction was shown to compromise -cell secretory function. Note that altered Myt3 translation did not affect glucose clearance in female mice under normal physiology or high-fat diet challenge. The underlying reason is not known.

There are a couple of other unresolved issues. First, because *Myt3* produces several different messages with different 5’ ends, and our manipulation only targets one of them, only mild translational changes were induced by our manipulation. Additionally, we induced Myt3 upregulation prior to HFD treatment. Thus, we do not know whether the observed β-cell defects after HFD induction arose from upregulated Myt3 pre-treatment or down-regulated Myt3 during treatment. Future studies to follow β-cell changes at different time points of HFD treatment could address this issue. Second, the mutant allele was present in all cells of the mice, including those in the brain regions that can impact endocrine secretion and energy homeostasis. Thus, we do not know if the defective glucose homeostasis is a compound effect of multiple organs. To this end, transplanting mutant islets from newly born mutant mice into wild-type mice and following the islet phenotypes could address this question.

## Methods

### Animal models and procedures

Mouse usage is approved by the Vanderbilt IACUC for Dr. Gu. Euthanasia follows the guidelines of AALAC. The *Myt3^F^* and *Pdx1^Cre^* mice were described previously (19). To derive the *Myt3^IM^* mice, Cas9 mRNA was co-injected with a guide RNA near the mutated sequence and a single-stranded DNA oligo spanning the mutated region (with a 12-base insertion) into the fertilized egg and implanted for mouse production. Note that the mutation creates a new Hind III restriction site, allowing a mutation screen with PCR, followed by Hind III digestion. After identifying the first-generation founders, they were backcrossed with CBA/BL6j mice for four generations. Heterozygous mice were then intercrossed to derive wild-type control and homozygous mutant mice for characterization.

### DNA construct, transfection, and luciferase assays

The cDNAs of RLuc and FLuc plus polyA signals from a vector from Addgene (#226464) were PCR amplified and cloned into a vector containing a bi-directional TetO enhancer/promoter (Addgene, #96963), producing general vector pYW1134. Note that during cloning, cloning site NotI was inserted at the 5’end of Fluc for later insertion of 5’ cDNA regulatory elements. The 5’ end of Myt3 T2RC was amplified via PCR using oligos with overhanging sequences that overlaps with that of pYW1134, on either side of NotI. Gibson assembly was then used to clone the 5’ sequence into the vector pYW1134, producing reporter pYW1377 (containing the WT 5’ sequence). To introduce the mutation, similar approve was used except the mutated 5’ sequence was synthesized, producing pYW1381.

For translational tests, pYW1377 or pYW1381 and pCMV-rTTA plasmids were co-transfected into HEK293 cells and cultures at 37 °C. 24 hours later, some samples were shifted into 42-43 °C for 3 hours. Cells were then collected for RLuc and Fluc assays using a Dual-luciferase kit from Promega. The ratios between FLuc/Rluc were used to measure the translational efficiency. Note that we tested conditions with/without Dox, measuring translation at high or low transcription levels.

### Intraperitoneal glucose tolerance test (IPGTT), plasma hormone assays, and high-fat diet (HFD) treatment

IPGTT followed a routine method. Mice were fasted overnight (∼16 hours). Glucose was injected at 2g/kg in mice without HFD challenge or at 1g/kg after HFD challenge, followed by blood glucose measurement via tail vein nip. To induce obesity, ∼5-week-old mice were fed HFD [VWR, 60% calories from fat (compare with 12% from fat in control diet)] for three to five months.

### Marker immunofluorescence (IF)

Antibodies used were: guinea pig anti-insulin (Dako, A0564), goat anti-Pdx1 (gift from Chris Wright of Vanderbilt), rabbit anti-Myt1 (this lab), rabbit anti-Ki67 (Abcam, RRID: AB_443209), rat anti-Myt3 (this lab). Secondary antibodies are all from Jackson ImmunoResearch: Alexa-Flour-488-donkey anti-rat (RRID:AB_2340683), Alexa-Flour-594-donkey anti-rabbit (RRID: AB_2340621), and Alexa-Flour-647-donkey anti-goat (RRID:AB_2340437). All antibodies were used at a 1:1000 dilution. All these antibodies were authenticated using mutant tissues.

### Islet preparation and secretion assays

Islet isolation uses collagenase Type IV perfusion followed by handpicking in RPMI1066 media (with antibiotics, 11 mM glucose, and 10% FBS) (19). After picking, the islets were allowed to recover in RPMI1066 media overnight before insulin secretion assays. Secretion assays were done in standard KRB solution with 2.8 mM glucose (G2.8), G20, or (G20 + 30 mM KCl) (G20K). For each stimulation, a 45-minute window was used. Total insulin was obtained with ethanol alcohol extraction.

### ScRNA-seq and Real-time RT-PCR

Freshly isolated islets were washed with Ca^2+^/Mg^2+^ free HBSS for 3 × 10 minutes. They were then dissociated into single cells with trypsin. InDrop-seq was then used for RNA sequencing (targeting 120 million reads), Novaseq 6000, Illumina (29,30). Each sample has islets from two males and two females. DropEst was used to preprocess scRNA-seq reads and to generate count matrices (31). Cells with low uniquely mapping reads (<500), low proportion of expressed genes (<100) or high proportion mitochondrial RNAs (>10%) were removed. Reads were normalized using UMI-filtered counts. Cell subpopulations were identified and visualized by UMAP using Seurat based on the first 30 principal components generated from the top 2000 highly variable genes (32,33). Differentially expressed genes (DEGs) between mutant and control β cells were identified by Seurat at the criteria of |log2 fold change|> 0.50 & FDR< 0.05. The Database for Annotation, Visualization and Integrated Discovery (DAVID) was used for functional clustering analysis (34).

### Statistical analysis

Students’ *t*-test was used for pairwise comparisons at single time points or paired genotypes. Two-way ANOVA was used to compare multiple groups of data points. A *p*-value of 0.05 or lower was considered significant.

### Data Accessibility and request for materials

Upon manuscript acceptance, the RNAseq data will be available in the Gene Expression Omnibus (GEO). Further requests for resources and reagents should be directed to and will be fulfilled by.

## Author contributions

RH, MY, XT, and GG performed mouse physiological assays, islet picking/secretion assays, IF staining, and imaging. YW and QL analyzed scRNA-seq data. YX, AJS, and KSL performed scRNA-seq using InDrop. KL, QL, and GG conceptualized the study. All authors participated in manuscript writing/proofing.

