## Supplemental Figures for "Deregulated Myt3 translation predisposes islet β-cells to dysfunction under obesity-induced metabolic stress"

**Summary of Supplementary Materials:**

1) Materials and Methods.
2) Supplementary table description (1 table).
3) Supplementary Figures and legend (three figures).

**Materials and Methods:**

IPGTT, GISS assays, RNA-seq data analysis, and IF analysis utilized methods described in the manuscript.

**Supplementary Table S1.** Spreadsheet for gene expression analysis.

### Supplementary Figures:

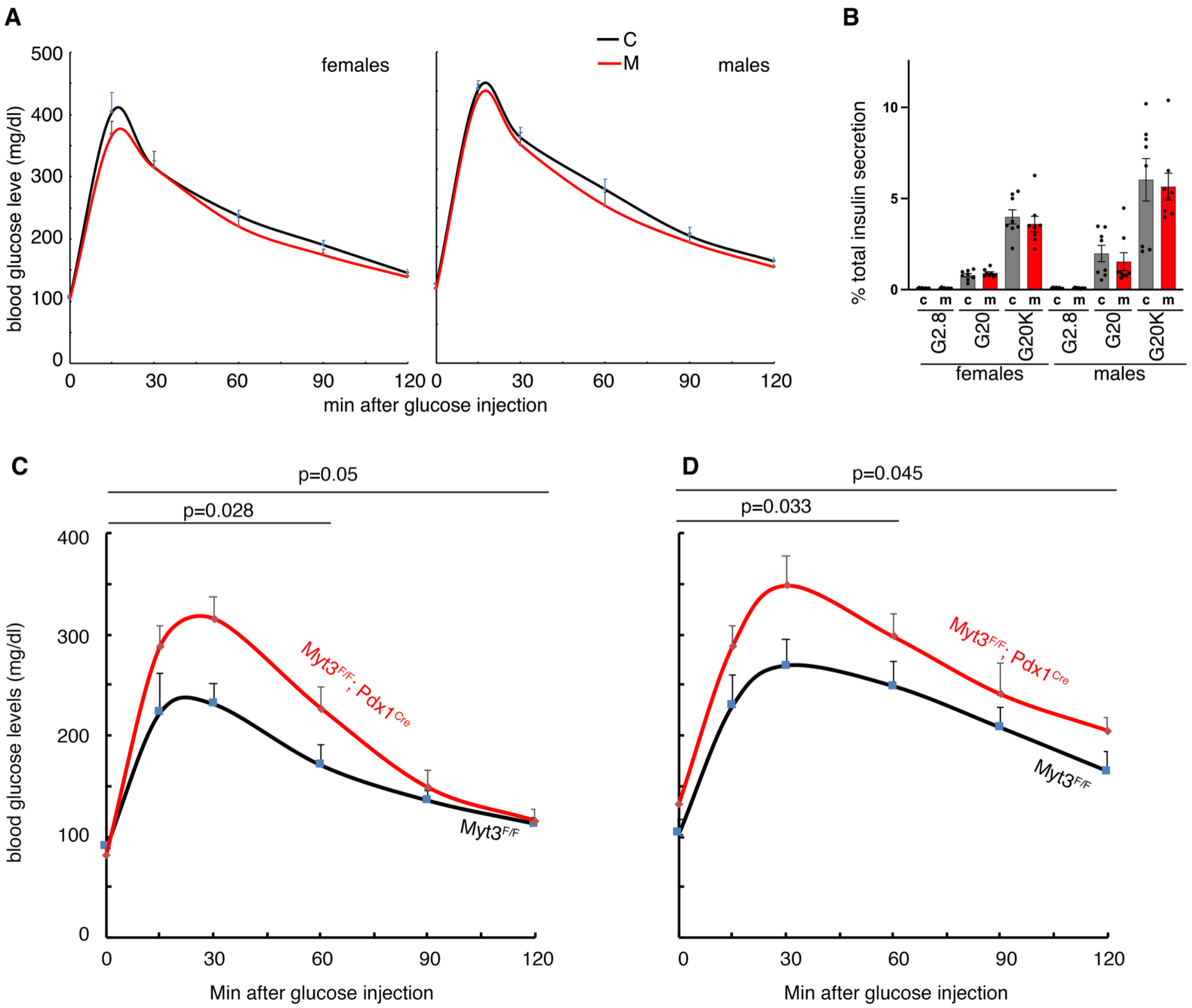


**Figure S1. Myt3 is required for glucose homeostasis in aged mice.** (A) IPGTT of 8-week-old mice. In both sexes, 6 controls and 6 mutants were tested. (B) Islet GSIS of 8-week-old mice, shown as % of total insulin secreted within a 45-minute window. Each assay contained at least 4 mice, 2-3 technical repeats from each mouse. (C, D) IPGTT of ~5-month-old mice. In females, 6 controls and 6 mutants were tested. In males, 6 controls and 7 mutants were tested.

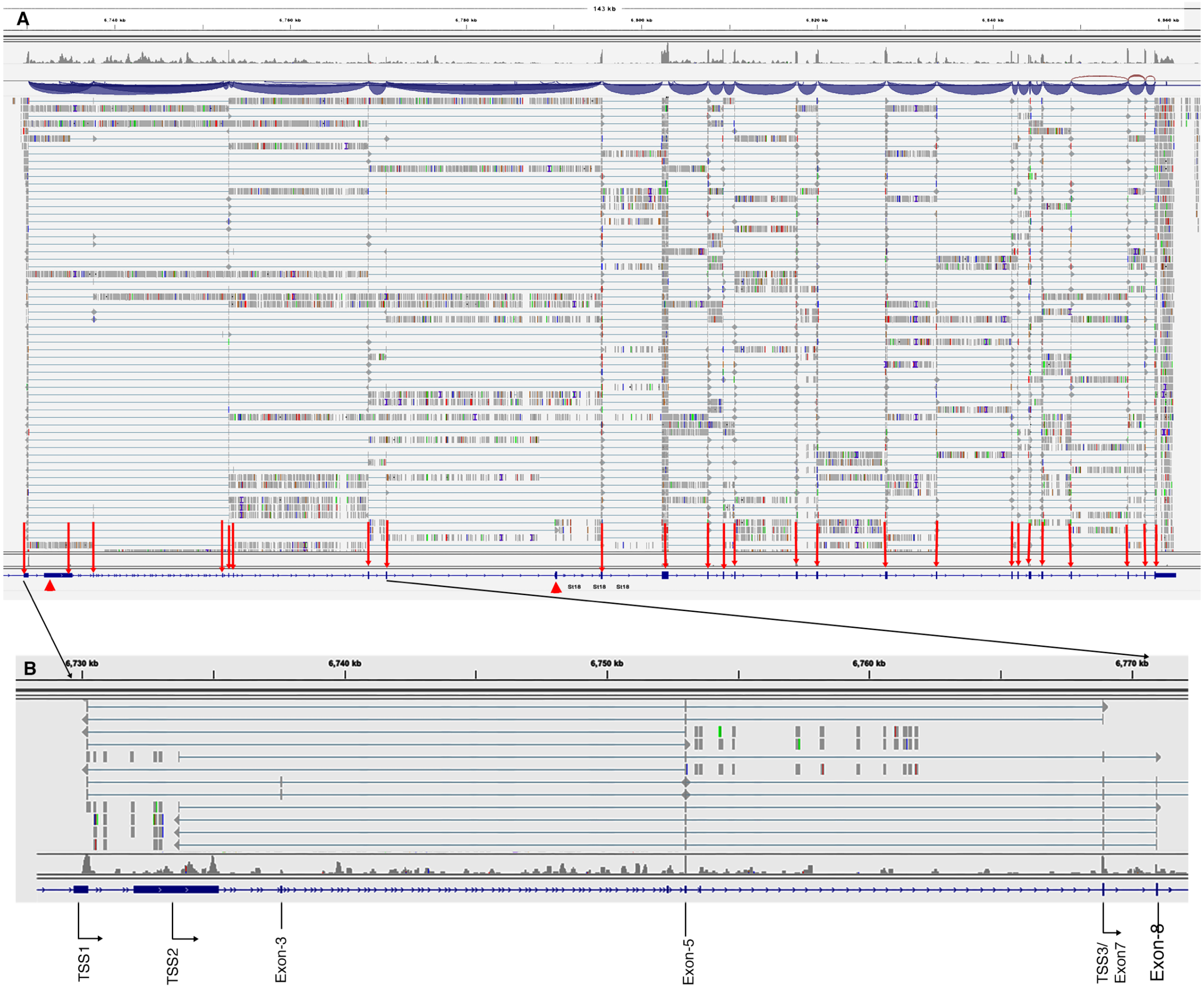


**Figure S2. Exon expression of Myt3 in purified adult β cells.** The original RNAseq data will be posted upon publication.

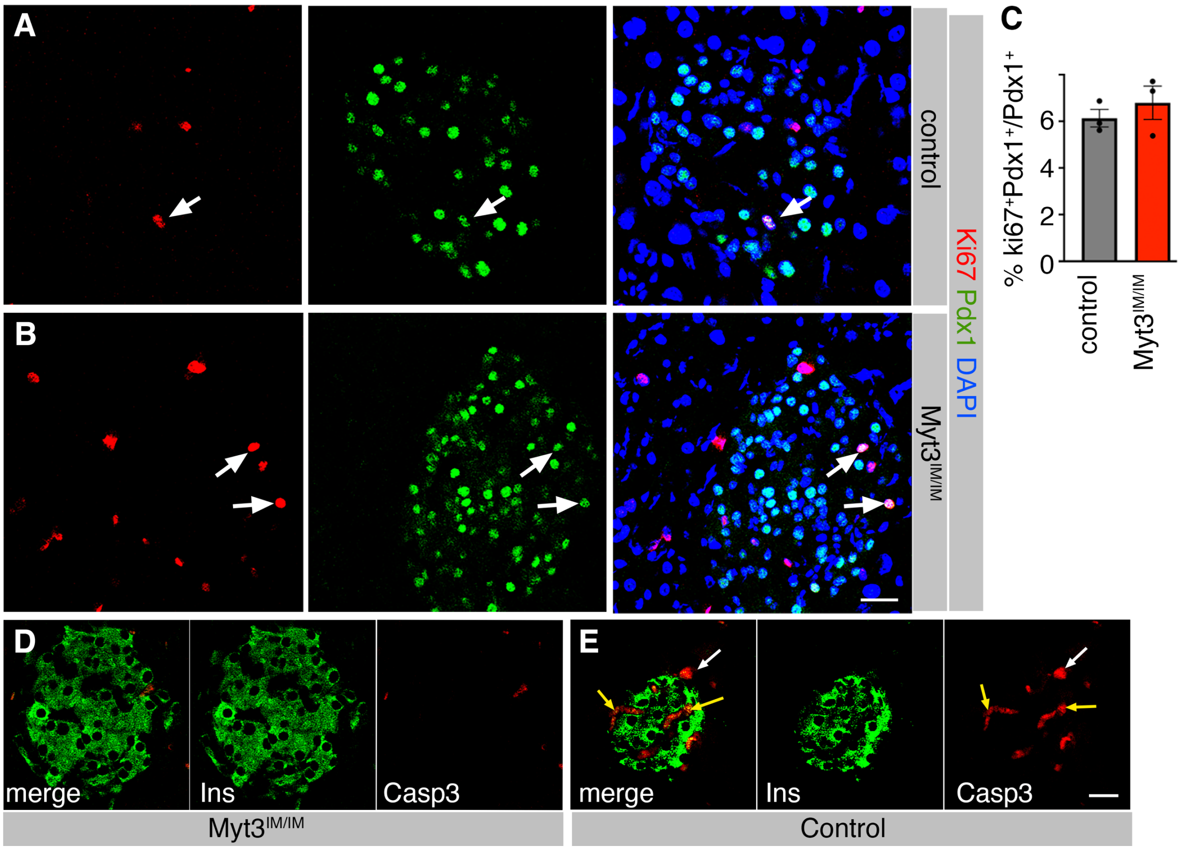


**Figure S3. Myt3$^{IM/IM}$ cells have normal proliferation and viability under HFD treatment.** (A-C) Ki67 and Pdx1 staining in control and Myt3$^{IM/IM}$ islet cells after 3-month feeding with HFD. The white arrow, a Ki67$^{+}$Pdx1$^{+}$ cell, appears to be dividing. (D, E) Apoptosis assays (via cleaved caspase 3, Casp3) in β cells of control and Myt3$^{IM/IM}$ islet cells after 3-month feeding with HFD. White arrows, a Casp3$^{+}$Ins$^{-}$ cell, to show that the IF assay was working. Yellow arrows, blood cells, recognized by their shape and location within the islets. Bars, 20 μm.
